# Opportunistically collected photographs can be used to estimate large-scale phenological trends

**DOI:** 10.1101/794396

**Authors:** Shawn D. Taylor, Robert P. Guralnick

## Abstract

**Premise:** Research on large-scale patterns of phenology have utilized multiple sources of data to analyze the timing of events such as flowering, fruiting, and leaf out. In-situ observations from standardized surveys are ideal, but remain spatially sparse. Herbarium records and phenology-focused citizen science programs provide a source of historic data and spatial replication, but the sample sizes for any one season are still relatively low. A novel and rapidly growing source of broad-scale phenology data are photographs from the iNaturalist platform, but methods utilizing these data must generalize to a range of different species with varying season lengths and occurring across heterogenous areas. They must also be robust to different sample sizes and potential biases toward well travelled areas such as roads and towns.

**Methods/Results:** We developed a spatially explicit model, the Weibull Grid, to estimate flowering onset across large-scales, and utilized a simulation framework to test the approach using different phenology and sampling scenarios. We found that the model is ideal when the underlying phenology is non-linear across space. We then use the Weibull Grid model to estimate flowering onset of two species using iNaturalist photographs, and compare those estimates with independent observations of greenup from the Phenocam network. The Weibull Grid model estimate consistently aligned with Phenocam greenup across four years and broad latitudes.

**Conclusion:** iNaturalist observations can considerably increase the amount of phenology observations and also provide needed spatial coverage. We showed here they can accurately describe large-scale trends as long as phenological and sampling processes are considered.

## INTRODUCTION

Earlier flowering and leaf onset in the spring is one of the strongest indicators of climate change (Parmesan and Yohe, 2003; Scheffers et al., 2016). Tracking and forecasting these trends for the myriad of plant species remains challenging due to the need for a large amount phenological observations across a species range (Chuine and Régnière, 2017). Large-scale observing networks and preserved herbarium specimens have provided a wealth of information, yet still leave large areas under-represented. Here we explore a new data stream of phenological observations from iNaturalist, an online social platform where users submit photos of plants and animals annotated with the time and geographic coordinates. Other users submit identifications of the taxon, resulting in a consensus vote on the final species determination. Observations of plants on the platform commonly include flowers in the photo, thus can indicate the time and location a species was in flower and is analogous to imaged herbarium records now commonly made available via aggregators such as iDigBio. With an adequate sample size over time and space, the flowering phenology can potentially be inferred for the entire growing season across a large portion of a species range.

There is no standardized method yet developed to model large-scale phenological trends across space from dispersed phenological observations. Potential methods utilizing these data need to account for several confounding factors, as the observations are derived from the interaction of both spatially heterogeneous phenological patterns and sampling bias. At a single location the full seasonal phenology of a species can resemble various statistical distributions such as the normal, beta, or uniform distribution. As the spatial extent expands the full phenological distribution can change depending on sample size and location (Fig. 1) (de Keyzer et al., 2017). Observations pull from this underlying phenology, but may have their own spatial and temporal observer bias. As with other citizen science projects, we expect iNaturalist observations to have spatial bias toward roads and populated areas (Dickinson et al., 2012). Similar to herbarium records we also anticipate a temporal bias toward in-season observations of flower presence (Daru et al., 2018). Thus, a method to estimate onset at large spatial scales should be generalizable to a variety of phenologies and also accommodate highly clustered sampling with little to no flowering absence represented.

**Figure 1:**
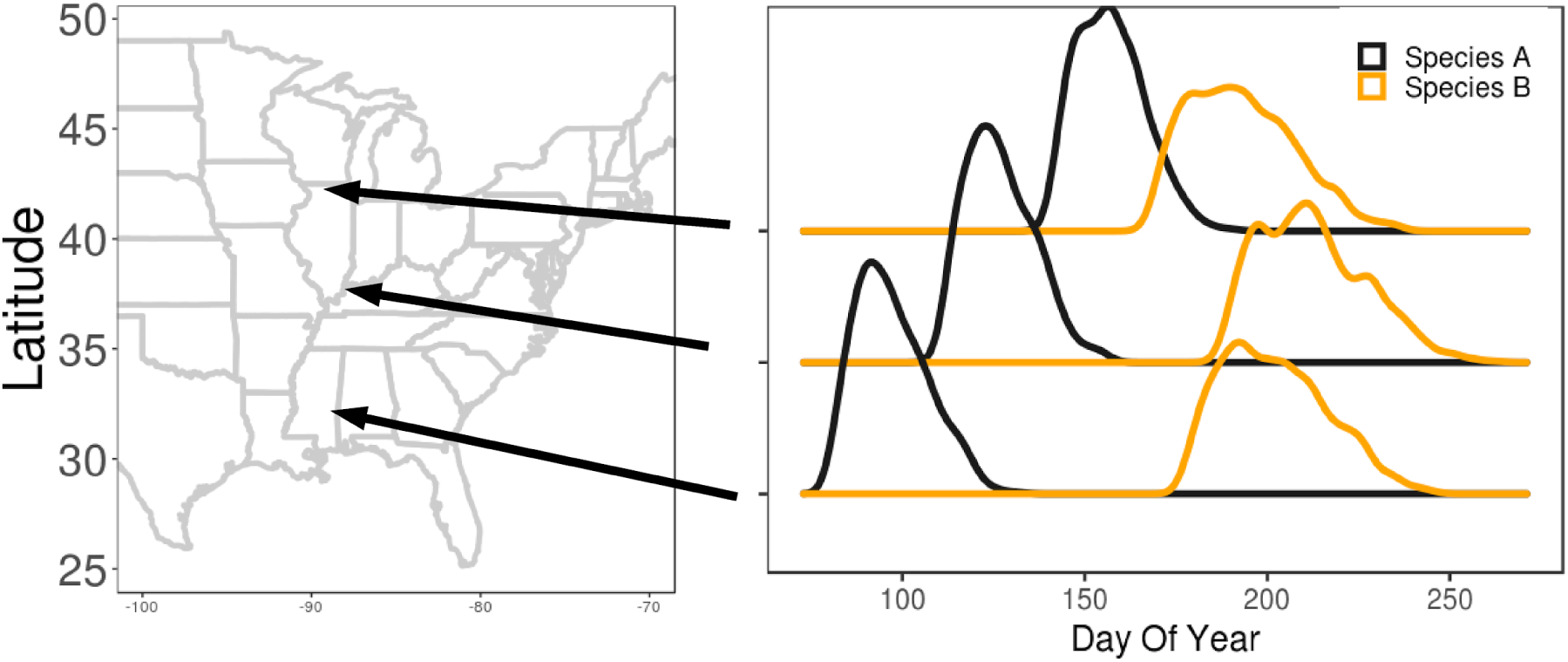
The flowering distribution of two theoretical plant species along a latitudinal gradient. Species A shifts linearly with increasing latitude while Species B shifts non-linearly.

In this study we developed a method to estimate the onset of flowering across large-spatial scales using a well-tested phenological estimator based on the Weibull distribution. Using the Weibull distribution is a well tested method and is beneficial as it utilizes only flowering presence data (Pearse et al., 2017; Taylor, 2019). We then integrated the Weibull estimator into a spatial ensemble framework (Fink et al., 2010). In the first part of our analysis, we test the method using simulated data where the underlying phenology and sampling scenarios are known. In the second part, we apply the methods to actual iNaturalist data and compare the estimates of flowering onset to landscape level greenup from the Phenocam network. Finally we discuss the potential for broadest use of iNaturalist data in ecological and phenological research.

## MATERIALS AND METHODS

### The Model

Our model, the Weibull Grid model, combines a Weibull distribution based estimator with a spatially explicit ensemble to estimate flowering onset across a heterogeneous landscape. The Weibull distribution can model a large range of shapes, and is commonly used to used to estimate the start or end of a process (Roberts and Solow, 2003; Pearse et al., 2017). It utilizes only observations of flowering presence, thus is advantageous here since iNaturalist data has a potential in-season biases. The estimated date of first flowering is the sum of the dates of all flowering weighted by the joint Weibull distribution and is equivalent to estimating an origination or extinction date. Taylor (2019) found it to be the superior estimator for flowering onset in most cases, and at least as good as methods that also include sparse absence reporting.

To use the Weibull estimator across heterogeneous landscapes we incorporated it into the random spatial grid used by Fink et al. (2010). A landscape is first divided into equally spaced grids. Within each grid cell boxes are uniformly randomly located. Observations within each box are then used in the Weibull estimator to estimate onset, thus all boxes have a spatially independent estimate. An estimate of onset at any point on the landscape is the median estimate from all boxes in which the point resides, with quantiles of the estimates used for confidence intervals. Model parameters are 1) the sizing of the initial grid cells, 2) the sizing of the boxes, and 3) the number of boxes per grid cell.

### Simulated Data

To evaluate the Weibull Grid model we first used simulated flowering data to create scenarios with a variety of underlying phenology and sampling schemes. We first simulate flowering on an x,y coordinate system designed to represent longitude and latitude, respectively. We used four parameters to describe the underlying phenology: 1) the initial start day of year (DOY) of flowering, 2) the length of flowering, 3) whether the spatial gradient is linear (ie. the change in the start DOY is a function of only the y-axis) or nonlinear (ie. a random surface is generated over which onset varies, Fig. S1), and 4) the strength of the spatial gradient. The start DOY is adjusted across the landscape by the equation ‘DOY ∼ initial DOY + y*strength’ when the gradient is linear, and ‘DOY = initial DOY + β*strength’ when the gradient is non-linear, where β is a random surface scaled to 0-1. The random surface is generated with the R package gstat using a variogram with a Gaussian distribution, a nugget of 0, sill of 0.025, and range of 0.6.

The true flowering curve is a normal distribution with the mean DOY at the midpoint of flowering, and the start DOY and the end doy (start DOY + flowering length) each at three standard deviations. From this, observations of flowering are randomly drawn using the distribution probabilities as weights. Simulated observations of flowering are randomly drawn from all DOYs 1:365 where peak flowering (the center of the distribution) has the highest probability, and any day before the Start DOY or after the End DOY has 0 probability.

The location of each observation in space is simulated to represent completely random sampling (non-clustered), or bias sampling toward 1 to several locations (clustered). Non-clustered sampling was generated by randomly selecting x,y coordinates from uniform distributions. Clustered sampling was generated using a Thomas process, where a random number of cluster centers was chosen from a Poisson distribution (λ=5) and n samples are generated with the highest probabilities centered on the cluster centers. Finally the total sample size of flowering observations was simulated to represent both abundant and relatively rare species.

We used a range of parameters to represent the potential phenology of as many species as possible. We tested flowering lengths of 15, 30, 45, and 60 days (Table 1). Using satellite data from a prior study we found that spring greenup in the Eastern U.S.A. had a trend of 3.36 days latitude^−1^ (Melaas et al., 2018). From this we derived three potential gradient strengths labeled Weak (1.68), Moderate (3.36), and Strong (6.72). In the simulation study the extent is 0-1, thus the gradient strengths were scaled to 0.1 so 10 degrees of latitude is simulated. Finally the spatial gradient type could be either linear or nonlinear. For the sampling scenarios we used sample sizes of 150, 300, 600, and 1200 flowering observations, where the observations could be uniformly randomly distributed or clustered. For each combination of parameters from both the underlying phenology and sampling scenarios we ran 100 simulations, with 19,200 in total.

**Table 1:** Parameters used in the simulation study.

| Parameter Category | Parameter | Range of Values | Description |
| --- | --- | --- | --- |
| Underlying Phenology | Flowering Length | 15,30,45,60 | The length of the flowering period. |
| Underlying Phenology | Gradient Strength | 1.68, 3.36, 6.72 | The strength of the spatial gradient |
| Underlying Phenology | Gradient Linearity | Linear or Nonlinear | Whether the gradient changes only w/ respect to the y-axis, or a random surface. |
| Sampling Scenario | Sample Size | 150, 300, 600, 1200 | The number of observations |
| Sampling Scenario | Observation Clustering | Clustered or Nonclustered | Whether observations are spatially clustered or random. |
| Weibull Grid Model | Box Size | 0.2, 0.4 | The size of the randomly distributed boxes. |
| Weibull Grid Model | Number of boxes | 5, 10, 20, 40 | The number of randomly located boxes within each grid cell |
| Weibull Grid Model | Grid Cell Size | 0.1, 0.2, 0.3 | The cell size of the initial grid. |

Each simulation was fit with the Weibull Grid model several times using different combinations of parameters (Table 1). We tested initial grid cell sizes of 0.1, 0.2, and 0.3, box sizes of 0.2 and 0.4, and boxes per grid cell of 5, 10, and 20, 40. For each simulation we also fit a Naive model to test the performance of the Weibull Grid model. The Naive model used a linear regression of DOY∼y, where y, representing latitude, is assumed to be the primary spatial gradient along which phenology varies. The Naive model onset estimate is then 0.01 percentile prediction interval.

For each simulation we generated 441 evenly spaced points on the simulated grid representing the true flowering onset date and calculated the RMSE of the models estimated onset date at the same locations. For each combination of Weibull Grid parameters, underlying phenology, and sampling scenarios we calculated, using the 100 replicate simulations, the average: 1) RMSE, 2) bias, and 3) percentage of the 441 testing observations which had no estimates. A location can lack an estimate in low sample size scenarios if none of its associated boxes had the minimum sample size (n=3) needed to produce an estimate. For each phenology and sampling scenario, we chose the set of Weibull Grid model parameters with the lowest RMSE to present here. The lowest errors for all simulated scenarios and the corresponding Weibull Grid parameters are available in the appendix (Table S1, Fig S2-S3).

### iNaturalist and Phenocam comparison

We compared estimates of onset using iNaturalist flowering data for two species and compared them with greenup estimates from Phenocam sites. We downloaded all research grade observations of *Rudbeckia hirta* and *Maienthemum canadense* from the dates 2016-01-01 to 2019-08-01 (GBIF.org, 2019). From the primary photo for each observation we scored whether a fully open flower was present or not. We scored *R. hirta* as “flowering” when at least 1 flowering head with at least 50% of visible ray petals were fully unfolded and non-senesced. We scored *M. canadense* as “flowering” when at least 1 fully open flower with non-senesced petals was visible. We kept all observations scored as “flowering” and with coordinates accurate to at least 50km. We built Weibull grid models to estimate flowering onset for each species and year (8 onset models total for the 2 species across 4 years). Weibull Grid parameters were chosen from the optimal parameters from the simulation analysis described above, and were based on the sample size, range size, and flowering length for each species (Figs. S2-S3).

We matched Phenocam locations which best represented the primary habitat and range for the two species, and which had data available in the years 2016-2019. For *R. hirta* we chose 6 grassland sites in the continental U.S.A. east of longitude -100. For *M. canadense* we chose 6 deciduous broadleaf sites in the N.E. U.S.A., Great Lakes Region, and southern Appalachian Mountains (Table S2). Using the phenocamr package we downloaded and extracted greenup estimated by the 10% rising transition of the 3-day 90th percentile Gcc and the associated confidence intervals (Richardson et al., 2018). For each Phenocam site location the flowering onset from the respective Weibull Grid model (*R. hirta* models for phenocam grassland sites, and *M. canadense* models for phenocam deciduous broadleaf sites, matched to the respective year) was estimated for comparison with the phenocam derived greenup date.

All analysis was done using the R programming language (version 3.6.0, R Core Team 2017). Packages used during the analysis included phest (Pearse et al., 2017), spatstat (Baddeley et al., 2015), gstat (Gräler et al., 2016), doParallel (Weston and Calaway, 2019), dplyr (Wickham et al., 2017), tidyr (Wickham and Henry, 2018), ggplot2 (Wickham, 2016), testthat (Wickham, 2011), and lubridate (Grolemund and Wickham, 2011).

## RESULTS

### Simulation Analysis

Using simulated flowering observations with known onset dates the Weibull Grid model performed best when the underlying phenology had a non-linear spatial gradient (Fig. 2). When the underlying phenology of the simulated flowering data was based on a linear spatial gradient, the Naive model outperformed the Weibull Grid model in nearly all cases (Fig. 2, Table S1). When the underlying phenology was a non-linear spatial gradient, the Weibull Grid model outperformed the Naive model, except with long flower days of weak and moderate gradients (Fig. 2 B,D). Performance of both models generally decreased (ie. had higher error) with increasing flowering length, though with a strong spatial gradient the flowering length had either no effect on the Weibull Grid model (Fig. 3 F), or had the highest errors at the shortest and longest flowering lengths (Fig. 3 E).

**Figure 2:**
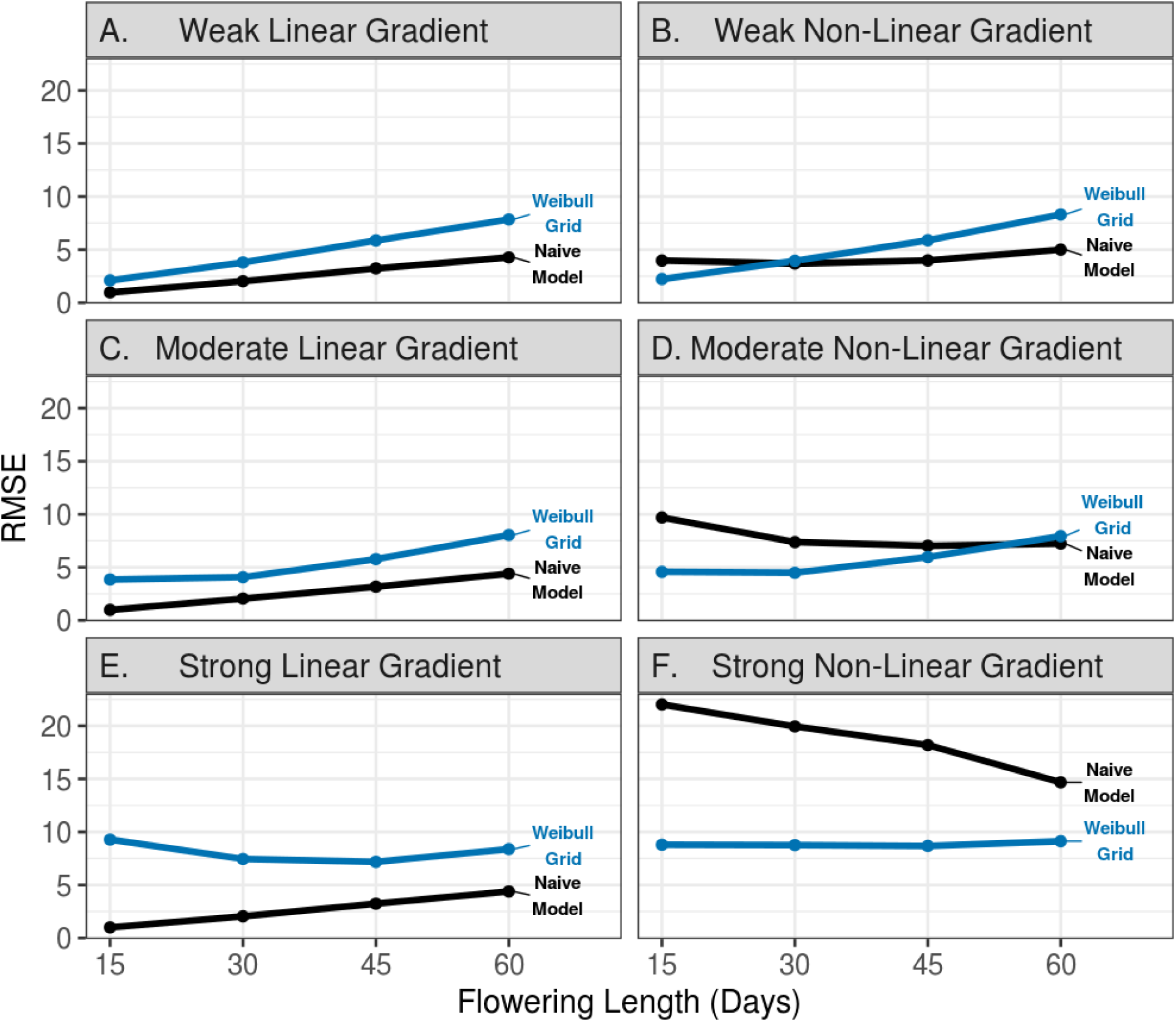
The RMSE of the Weibull Grid model and the Naive model in estimating onset for 6 types of flowering gradients (A-F) and 4 flowering season lengths (x-axis), and using a sample size of 300 based on clustered sampling. Each point represents the average RMSE from 100 simulations where the true onset date was specified.

**Figure 3:**
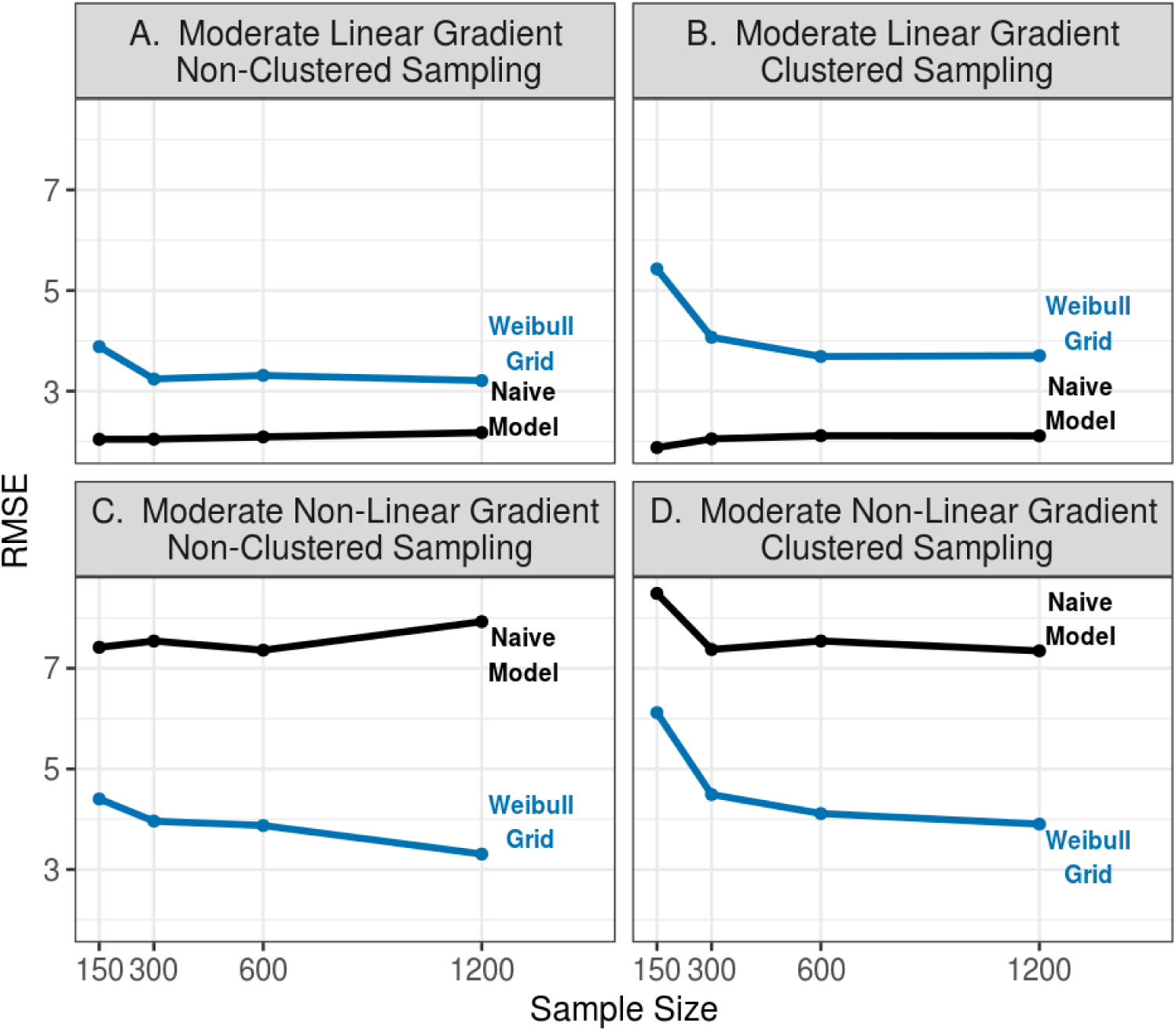
RMSE of the Weibull Grid model and a Naive model in estimating onset from for either non-clustered (A-B) or clustered sampling (C-D). The underlying phenology is a moderate spatial gradient with a length of 30 days and spatial gradient linearity specified. Each point represents the average RMSE from 100 simulations where the true onset date was specified.

Simulating clustered sampling, as opposed to completely random sampling, increased errors for the Weibull Grid model regardless of the underlying phenology (Fig. 3, Table S1). The change in error was highest when the sample size was the lowest. Using a sample size of 150 simulated observations the Weibull Grid model RMSE increased by more than 1 day when clustered sampling was introduced. This increase happened regardless of flowering length or the spatial gradient strength (Table S1). As the sample size increased the magnitude of the RMSE change attributed to clustered sampling decreased. Higher sample sizes improved the Weibull Grid model performance overall, especially moving from 150 to 300 samples. Beyond 300 simulated samples performance improvements were relatively small.

### Comparison of iNaturalist based estimates with Phenocam estimates

We used the Weibull Grid model to estimate flowering onset for two species based on flower phenology observations derived from iNaturalist photos. We compared these estimates with greenup dates from the Phenocam network. In deciduous broadleaf forests the Weibull Grid model estimated flowering onset of *M. canadense* an average of 2.7 days after canopy greenup (range: −10.4 to 17.3). At grassland sites the Weibull Grid model estimated flowering onset of *R. hirta* after canopy greenup at all sites and years, 33.9 days on average (range: 1.3 to 66.1). However, some estimates, especially at lower latitude sites, had confidence intervals extending well before the Phenocam greenup.

## DISCUSSION

Here we have shown that opportunistically collected photos of in-flower plants from iNaturalist can provide consistent estimates of large-scale phenological patterns. We showed that a method to estimate flowering onset integrated into a spatial ensemble framework can consistently match independent estimates of greenup. The same method was also validated using simulated data and was especially beneficial when non-linear spatial gradients were present. The iNaturalist platform is still relatively new, and its data has largely been used as verified observations in species distribution models. Extracting ecologically relevant information from user submitted photographs is challenging (Barve et al., 2019), yet the scale of the data can provide an extremely high return on investment for phenological research.

### Diverse Data Harmonization

The sheer amount of observations available from iNaturalist as well as the spatial coverage provides potential for analysis at scales not before possible. The data density provides high replication across gradients, which provides the most information about large-scale patterns (Kreyling et al., 2018). Yet, as a relatively new platform which does not emphasize standardized sampling, iNaturalist data are missing historical context as well as site fidelity. Herbarium specimens have been a primary source of large-scale analysis and provide historic observations dating back over 100 years (Willis et al., 2017). Yet for the two species used here historic herbarium observations rarely exceed 100 total observations per year, thus are likely lacking adequate spatial replication to track seasonal large-scale trends (Fig. 5). Since 2016 the annual number of research grade *R. hirta* observations on iNaturalist has increased from 500 to nearly 2000. USA-NPN has similar increases in the number of observations, but does not provide as high a density since many observations are repeated at the same location (Fig. S4). Even with this lower density the repeated sampling observations from the USA-NPN, and other monitoring networks, helps constrain phenological timings (Gerst et al., 2016; Elmendorf et al., 2019). Thus with its dense sampling iNaturalist derived phenological data can compliment as opposed to replace other data sources.

**Figure 4:**
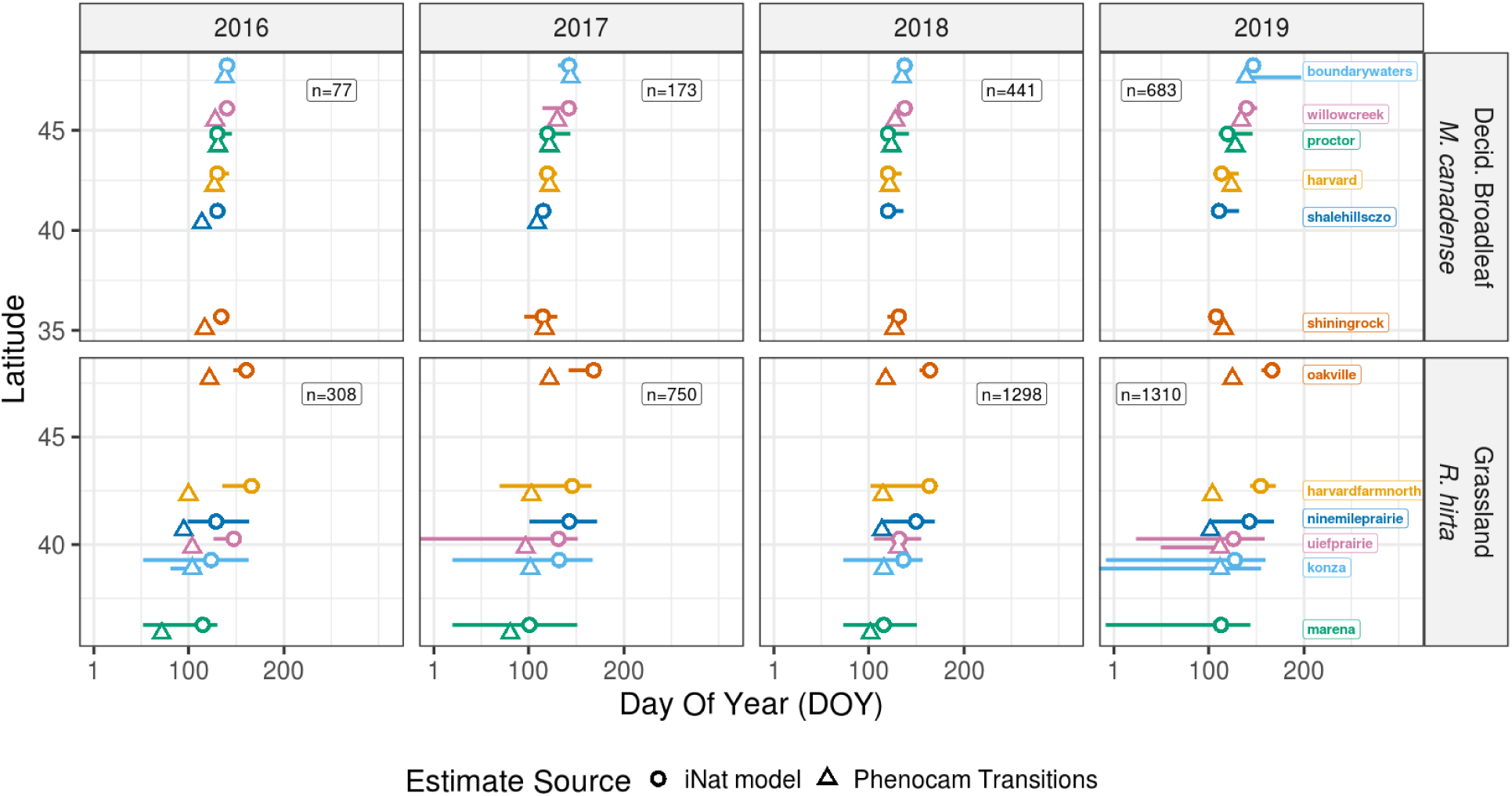
Flowering onset estimated from iNaturalist phenology data using the Weibull grid model (circles) compared with Phenocam derived estimates of canopy greenup (triangles). Colors represent Phenocam sites. Lines represent the 95% confidence interval, which may be not be visible if the interval is extremely small. Note that sites marena in 2019, ninemileprairie in 2017, and shalehillsczo in 2018-2019 do not have Phenocam estimates.

**Figure 5:**
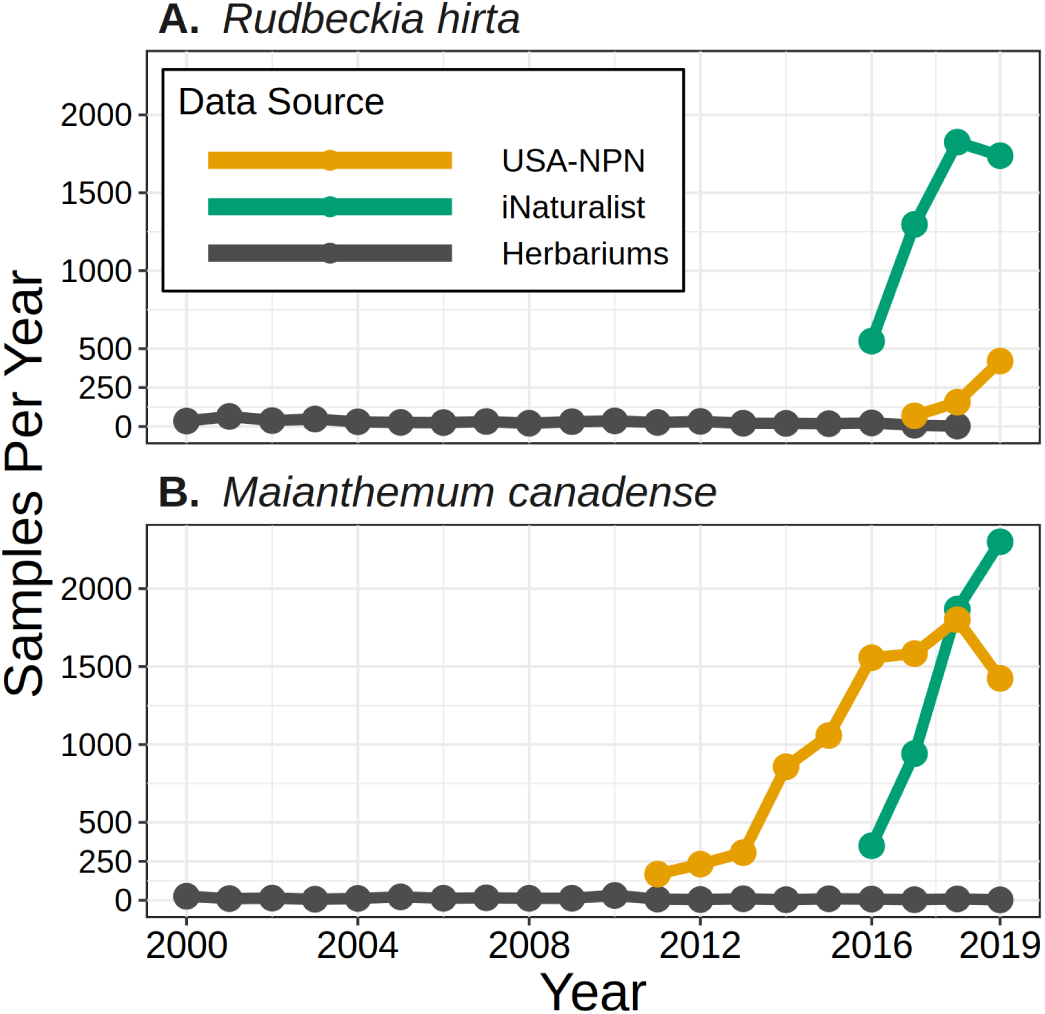
Total observations from other large-scale phenological data sources for the two species (A & B) analyzed in the current study. iNaturalist data reflects the total research grade observations regardless of flower status. USA-NPN data reflects observations of open flowers (phenophase 501, either present or absent) from Jan. 1, 2009 to Aug. 1, 2019 (USA National Phenology Network 2019). Herbarium data were obtained from iDigBio and reflects all observations marked as preserved specimens for the respective species. Though the x-axis is cropped from the year 2000 onward, herbarium observations here date back to the year 1800.

Combining these diverse data streams will be a major challenge, as they have a variety of protocols and observation biases. The Plant Phenology Ontology (PPO) provides a framework for harmonizing different phenology observations into common indicators (Stucky et al., 2018; Barve et al., 2019; Brenskelle et al., 2019). Hierarchical models can be used to model the different observations processes under a common underlying phenology (Ogle et al., 2015; Elmendorf et al., 2019) and also share strength between closely related species or ones with similar traits (Theobald et al., 2017). Incorporating potential drivers into models may also be difficult. For example if gridded temperature data were combined with onset estimates from the Weibull Grid model then the spatial grain chosen will affect the results (Jelinski and Wu, 1996). Alternatively, the presence/absence observations could be integrated into a model directly without first estimating onset dates (Clark et al., 2014; Elmendorf et al., 2019). Future analysis using the strengths of each dataset will likely provide the most information for inferring phenological patterns and their drivers.

### Weibull Grid Model Performance

The Weibull Grid model was most beneficial when the underlying phenology had a non-linear spatial gradient. This is due to the spatial ensemble method having a location dependent estimate which is informed only by nearby observations (Fink et al., 2010). This limits the model to regions with dense sampling, which we expect iNaturalist to provide for many abundant well-known species (Fig. 5). Estimates improved up to 300 total observations, but beyond that improvement of overall performance was minimal. Our analysis simulated 10 degrees of latitude/longitude, thus as a first approximation we can suggest, for every year of transitions to estimate, sampling densities of 3 observations per degree^−2^ at mid-latitudes. When the underlying phenology had a linear spatial gradient the Naive model always performed better. This scenario will likely be encountered when an analysis has a small spatial extent, and the onset of flowering, or other phenophases, is nearly simultaneous across the study region.

The Weibull Grid model was fairly robust to clustered sampling, with errors increasing by 1 day or less when clustered sampling was introduced and sample size was 300 or greater (Fig. 3). The model performed best with shorter flowering seasons and moderate to strong nonlinear spatial gradients. As the flowering season was lengthened observations on average were farther from the true onset date, thus the estimate had higher uncertainty. With stronger gradients, nearby observations were also more likely to be farther apart in time, again contributing to higher uncertainty and overall error. Larger sample sizes did improve performance to a degree, especially at the strongest gradient (Table S3).

We found a larger box size (sized 0.4 on a 0-1 simulated grid) had the lowest errors except in two cases: With a strong non-linear gradient combined with a short flowering season, and with a high sample size (n=1200). (Figs. S2-S3). The optimal choice for the other two parameters (the initial grid cell size and number of boxes within each cell) varied as the interaction between these two parameters determined the total number of boxes across the entire simulated landscape. Since the final point estimates are based off quartiles, more estimates from a higher amount of boxes are not detrimental and the only downside is increased computational time. Thus a small initial grid cell size and high number of boxes is recommended.

### iNaturalist vs. Phenocam Derived Onset Estimates

The flowering onset estimates for *M. canadense* and *R. hirta* were consistent with ecosystem level estimates. We had no a priori assumptions about when in the growing season the two species flowered, yet the Weibull Grid model fit with iNaturalist derived phenology data consistently provided estimates near Phenocam derived greenup. The onset of flowering for *R. hirta* was always estimated after Phenocam greenup, while *M. canadense* was estimated to be fairly close to the Phenocam greenup.

Estimates for *R. hirta* flowering onset had wider confidence intervals overall. With a larger range, and thus stronger gradient, and longer flowering season of *R. hirta* onset is inherently harder to estimate. At lower latitudes open flowers were observed in every month of the year. Recent direct measurements for *R. hirta* flowering onset are DOY’s 169-177 at 46.9 latitude (Dunnell and Travers, 2011), which is similar to the estimates of 160-166 at the oakville site.

Our results show that *M. canadense* flowers near canopy greenup throughout its range. *M. canadense* flowers in “early spring” (Flora of North America Editorial Committee 1993), and Helenurm and Barrett (1987) measured flowering onset between DOY’s 151-165 in Doaktown, New Brunswick (46.5 latitude) in the years 1979-1980. The Phenocam site willowcreek, at latitude 45.8, had *M. canadense* flowering onset estimates between DOY’s 138-142. This disparity could be due to the nearly four decades since the data collection by Helenurm and Barrett (1987), as another study showed *M. canadense* flowering 7 days earlier since 1980 due to warmer temperatures.

### The Potential Information Content of Photographs

We are discretizing the iNaturalist photos into binary presence or absence of open flowers, yet there is additional information in the photos which can be utilized. Phenophases such as fruiting, leaf onset and coloring can also be scored. The total number of flowers or fruit on an individual or in a photo can be tallied to quantify flower intensity or reproductive success. The natural progression of different phenophases can be used to constrain the timing of earlier and later transitions (Wolkovich and Ettinger, 2014; Ettinger et al., 2018). For example, fruiting follows flowering, thus the presence of fruit implies flowering onset happened at an earlier date, and vice versa. *M. canadense*, analysed here, had very discrete phenophase timing. During scoring we noted that open flowers of *M. canadense* never coincided with ripe fruit, but occasionally coincided with enlarging ovaries. We hypothesize the overlap between phenophases is highly species dependent and thus difficult to generalize in models. For example, some species have ripe fruit which persist for months, and the presence of maturing or unripe fruit may be a better phenophase to utilize. Likewise, long flowering plants with indeterminate inflorescences could simultaneously have unopened flower buds, open flowers, and immature and ripe fruit.

Other information beyond the presence or quantity of phenophases can be extracted from photos as well. Plants besides the target species may be present, allowing for analysis of competing species or community scale phenology. However, researchers should ensure the same photo was not submitted repeatedly to record multiple species. If the photo frame is large enough then landscape level greenness indices can be extracted using either machine learning and/or crowdsourcing techniques (Kosmala et al., 2016), which can be integrated with Phenocam data. iNaturalist observations identified as plant pollinators have a high chance of containing flowers, providing an avenue to analyse these phenological interactions (Gazdic and Groom, 2019). The long-term storage of iNaturalist photos and their metadata, similar to storing herbarium specimens, will provide future researchers with these analysis avenues, as well as other options which may not be obvious today (Heberling and Isaac, 2018).

### Limits and Biases of Phenology From Photographs

While we utilized only flowering presence data here, the absence of flowers or other phenophases can be highly informative in constraining estimates of phenological timings (Taylor, 2019). The scoring of iNaturalist photos provides a method for characterizing phenophase absence, though depending on size, abundance, and other traits, such absence reporting may be better suited to some species more than others.

We expected a bias toward observations of plants in flower on iNaturalist, and that was the case with one of the two species analyzed. For *R. hirta* 94% of observations had open flowers, while *M. canadense* had 36%. We hypothesize the presence of flowers drives a higher proportion of research grade observations in two ways: 1) the ability to positively identify a plant to the species level, and 2) the prominence of a plant driving the motivation of an observer to photograph it. *M. canadense* is a common, sometimes dominant, understory plant in N.E. North America with distinctive parallel veined leaves. This allows for both more observations and easier identification when flowers are absent, thus more non-flowering observations. *R. hirta* is common throughout Eastern North America, but the early season basal rosette can resemble other species in the Asteraceae family, thus lacking flowers identification is difficult. Without a positive identification the observations will not be marked as “research grade”, excluding them from most downstream analysis. Alternatively, if *R. hirta* is in the understory and lacking flowers it may not be prominent enough for an observer to take a photo to begin with. If the former is the primary driver for the dearth of non-flowering *R. hirta* research grade observations, then there is potentially a large history of phenological absence data available on iNaturalist if more advanced identification methods could be applied.

Extracting phenological information from photos has several strengths and limitations. False positives for flowers are likely low since the presence of flowers is easy to detect. There is potential dispute over what constitutes “flowering”, as the inflorescence of some species may remain highly visible even after successful pollination and cessation of ovule viability. Flowering heads comprising numerous flowers can also remain viable over several days to weeks. For these issues the PPO can provide guidance (Stucky et al., 2018), and multiple observers resulting in a consensus vote on final phenology decreases scoring errors (Barve et al., 2019). The potential for false negative (flowers absent) is high, especially if the entire plant is not included in the photo frame. *R. hirta* is a perennial taprooted herb, so it’s likely the full above ground portion of the plant, or at least the full inflorescence, is in most photos. *M. canadense* has spreading rhizomes, so it would be impossible to judge from a single photograph whether all stems of an individual plant are present. Larger plants will likely have higher error rates from scoring, either from only a portion of an individual being photographed or the plant being far away in the image (Barve et al., 2019). Photos of plants with charismatic flowers, even small ones, may exclude the rest of the plant which would prohibit deriving any information besides flower presence.

Taken together these biases will limit which plant species can have their phenology scored and analyzed from online photographs on iNaturalist and other platforms. Most suitable will be common, abundant and easily identifiable species which are large enough to be conspicuous, but small enough such that the entire plant is included in most photos at a near enough distance to adequately score phenophases. Still unstudied is the effect of observer effort across space and time. To obtain an adequate sample size a plant must flower (or have other phenophases present) at the locations and times coinciding with observers. Plants more abundant in locations which are more likely to be visited, and which flower during times mostly likely to be visited will be better suited. Tall species such as trees, rare species, or those with inconspicuous flowers will likely not be suitable.

### How Predominant is Nonlinear Phenology?

In our simulation analysis we found the Weibull Grid method to be superior when the underlying phenology is non-linear across space. In the real world we hypothesis that large-scale (>100km in extent) phenology gradients will have non-linear trends in the majority of plant species. Complex topography, which affects phenology thru abiotic drivers such as temperature and sunlight, predominates much of the world’s land surface. Even in areas of uniform topography differences in weather patterns, community structure, and successional status can change the phenology such that a single explanatory variable such as latitude cannot adequately account for the variance (Melaas et al., 2016). Regions experiencing shifts in climate at faster rates than surrounding regions will also contribute to non-linearity (Ault et al., 2015). In the cases where a linear gradient exists the Weibull Grid method may still be beneficial when the species encompass a large range, and the shape of the phenological distribution varies over latitude. Future research should explore further the approximate threshold in non-linearity in which a simple naive model fails to capture the variation. We admit that beyond studies using gridded climate or remotely sensed data, this aspect of plant phenology is largely unstudied, and we believe iNaturalist data will provide valuable information in analysing large-scale patterns.

## CONCLUSIONS

The iNaturalist platform has been operational since 2008, yet it’s grown to be one of the most popular citizen science platforms, given its inclusive participatory nature, tools to help with identification, and how it balances ownership issues related to photographs, while also strongly supporting open data approaches. Here we have shown the potential to extract phenological info from iNaturalist photographs, and outlined how these data, with their high spatial replication, can complement other sources. Quantifying large-scale trends across space and time can validate past findings of experimental studies and provide new hypotheses for future ones. Location specific transition estimates can also be used to parameterize models for large-scale forecasts, which benefit from data across a large spatial extent (Taylor, 2019). We made a spatially explicit model which showed, given an adequate sample size, iNaturalist derived data can provide precise estimates of phenological onset for plant species which occur across heterogeneous environments. Flexible modelling approaches which can utilize the unique structure of these data, eg. the Weibull estimator, will be key in large scale analysis. In the future, and as the iNaturalist database grows, it will likely become a primary source of phenological data.

## Acknowledgments

This research was supported by the Gordon and Betty Moore Foundation’s Data-Driven Discovery Initiative through Grant GBMF4563 to Ethan P. White. RPG acknowledges funding on a Macrosystems grant, NSF 1703048, to University of Florida and University of Connecticut. We thank all the contributors to iNaturalist, the USA National Phenology Network, Nature’s Notebook, and iDigBio. We thank Michael Belitz for feedback on early discussions of the manuscript.

## Author Contributions

ST conceived the idea and performed the analysis. ST and RG wrote the manuscript.

## Data Accessibility Statement

Code to fully reproduce this analysis is available on GitHub (https://github.com/sdtaylor/phenology_gradients, https://github.com/sdtaylor/flowergirds) and archived on Zenodo (https://doi.org/10.5281/zenodo.3473015).

## Appendix

**Table S1:** Errors for all combinations of phenology and sampling scenarios. Each number represents the RMSE of the best performing Weibull Grid model (chosen from a suite of potential parameter combinations, see main text & Table 1). Numbers in parenthesis indicate the RMSE of the Naive model for the same scenario. Bold text indicates when the Weibull Grid model had a lower RMSE than the Naive model.

|  |  |  | Sample Size |  |  |  |
| --- | --- | --- | --- | --- | --- | --- |
| Spatial Gradient Type | Gradient Strength | Flowering Length | 150 | 300 | 600 | 1200 |
| Clustered Sampling |  |  |  |  |  |  |
| linear | Weak | 15 | 1.9 (0.8) | 1.6 (1) | 1.7 (1) | 1.5 (1) |
| linear | Weak | 30 | 4.1 (1.8) | 3.3 (2.1) | 2.3 (2.1) | 2 (2.2) |
| linear | Weak | 45 | 6.6 (3.1) | 5.4 (3.2) | 3.6 (3.1) | 2.6 (3.2) |
| linear | Weak | 60 | 9.5 (4.3) | 7.3 (4.3) | 4.9 (4.3) | 3.4 (4.4) |
| linear | Moderate | 15 | 3.4 (0.9) | 2.1 (1) | 1.7 (1) | 1.5 (1) |
| linear | Moderate | 30 | 3.9 (2) | 3.2 (2) | 3.3 (2.1) | 3.2 (2.2) |
| linear | Moderate | 45 | 6.3 (3.1) | 4.9 (3.1) | 3.9 (3.3) | 3.7 (3.3) |
| linear | Moderate | 60 | 8.4 (3.9) | 6.9 (4.1) | 4.8 (4.2) | 4 (4.3) |
| linear | Strong | 15 | 6.2 (0.9) | 3.7 (1) | 3.3 (1) | 3.5 (1) |
| linear | Strong | 30 | 6.7 (2) | 4.3 (2.1) | 3.5 (2.2) | 2.9 (2.2) |
| linear | Strong | 45 | 6.4 (3.2) | 6 (3.2) | 5.8 (3.3) | 4.6 (3.3) |
| linear | Strong | 60 | 8 (4.3) | 6.7 (4.4) | 6.4 (4.4) | 6.6 (4.4) |
| non-linear | Weak | 15 | 2.2 (4.2) | 2 (3.8) | 1.9 (4.1) | 1.6 (4.3) |
| non-linear | Weak | 30 | 4.2 (3.4) | 3.4 (3.6) | 2.5 (3.3) | 2.2 (3.4) |
| non-linear | Weak | 45 | 6.8 (3.8) | 5.3 (3.6) | 3.8 (3.7) | 2.7 (3.6) |
| non-linear | Weak | 60 | 9.5 (4.8) | 7.5 (4.5) | 5 (4.6) | 3.6 (4.7) |
| non-linear | Moderate | 15 | 3.9 (10.9) | 2.5 (11.2) | 2.2 (10.6) | 2 (9.6) |
| non-linear | Moderate | 30 | 4.4 (7.4) | 4 (7.5) | 3.9 (7.4) | 3.3 (7.9) |
| non-linear | Moderate | 45 | 6.5 (6.7) | 5.3 (6.8) | 4.2 (6.6) | 4.2 (6.8) |
| non-linear | Moderate | 60 | 8.7 (6.7) | 7.1 (6.7) | 5.2 (6.9) | 4.5 (6.4) |
| non-linear | Strong | 15 | 7.1 (25.6) | 4.6 (23.9) | 4.4 (24.9) | 4.4 (25.3) |
| non-linear | Strong | 30 | 7.7 (20.7) | 5 (22) | 4.2 (21.5) | 3.8 (19.9) |
| non-linear | Strong | 45 | 8 (19.2) | 7.3 (17.9) | 6.2 (17.3) | 5.1 (16.7) |
| non-linear | Strong | 60 | 9.3 (16.3) | 8 (15.8) | 8.3 (16.7) | 6.8 (15) |
| Non-Clustered Sampling |  |  |  |  |  |  |
| linear | Weak | 15 | 2.5 (0.9) | 2.1 (1) | 1.8 (1) | 1.9 (1) |
| linear | Weak | 30 | 5.1 (2) | 3.8 (2) | 3.1 (2.1) | 2.5 (2.2) |
| linear | Weak | 45 | 7.8 (3) | 5.9 (3.2) | 4.6 (3.3) | 3.8 (3.2) |
| linear | Weak | 60 | 11.5 (4.3) | 7.8 (4.3) | 6.2 (4.5) | 4.8 (4.4) |
| linear | Moderate | 15 | 4.9 (1) | 3.9 (1) | 3.2 (1) | 2 (1) |
| linear | Moderate | 30 | 5.4 (1.9) | 4.1 (2) | 3.7 (2.1) | 3.7 (2.1) |
| linear | Moderate | 45 | 8.1 (2.9) | 5.8 (3.2) | 4.7 (3.2) | 4.2 (3.3) |
| linear | Moderate | 60 | 9.8 (4) | 8 (4.4) | 6.4 (4.3) | 5.3 (4.4) |
| linear | Strong | 15 | 9.7 (0.9) | 9.3 (1) | 5.7 (1) | 3.9 (1.1) |
| linear | Strong | 30 | 9.4 (2) | 7.4 (2) | 6.2 (2.2) | 4.1 (2.2) |
| linear | Strong | 45 | 9.9 (3) | 7.2 (3.2) | 6.8 (3.4) | 6.3 (3.3) |
| linear | Strong | 60 | 11.1 (4.1) | 8.4 (4.4) | 7.3 (4.4) | 7 (4.5) |
| non-linear | Weak | 15 | 2.7 (4.1) | 2.2 (4) | 2.1 (4) | 1.9 (3.9) |
| non-linear | Weak | 30 | 5.2 (3.7) | 3.9 (3.7) | 3.2 (3.4) | 2.6 (3.4) |
| non-linear | Weak | 45 | 7.8 (4.1) | 5.9 (4) | 4.5 (4.1) | 3.9 (4.1) |
| non-linear | Weak | 60 | 9.8 (4.8) | 8.3 (5) | 6.3 (4.9) | 5 (4.9) |
| non-linear | Moderate | 15 | 4.9 (10) | 4.6 (9.7) | 3.3 (8.7) | 2.4 (9.2) |
| non-linear | Moderate | 30 | 6.1 (8.5) | 4.5 (7.4) | 4.1 (7.5) | 3.9 (7.3) |
| non-linear | Moderate | 45 | 8 (6.6) | 6 (7) | 5.1 (7.5) | 4.5 (6.6) |

Table S1: Errors for all combinations of phenology and sampling scenarios.
| Spatial Gradient Type | Gradient Strength | Flowering Length | Sample Size |  |  |  |
| --- | --- | --- | --- | --- | --- | --- |
|  |  |  | 150 | 300 | 600 | 1200 |
| non-linear | Moderate | 60 | 10.3 (7.2) | 7.9 (7.2) | **6.5 (7)** | **5.5 (7.1)** |
| non-linear | Strong | 15 | **10.4 (25.8)** | **8.8 (22)** | **6.1 (20.4)** | **5 (24.9)** |
| non-linear | Strong | 30 | **11.3 (19.9)** | **8.8 (19.9)** | **6.8 (17.9)** | **4.8 (18.3)** |
| non-linear | Strong | 45 | **11.3 (16.4)** | **8.7 (18.2)** | **8.2 (17)** | **6.5 (15.4)** |
| non-linear | Strong | 60 | **11.9 (15)** | **9.1 (14.7)** | **8.7 (17)** | **8.1 (13.8)** |

**Table S2:** Parameters used for the Weibull Grid model for the 2 species using iNaturalist derived phenology data.

| Species | Primary Vegetation | Box Size | Grid Cell Size | Number of Boxes | Latitude Bounds | Longitude Bounds |
| --- | --- | --- | --- | --- | --- | --- |
| R. hirta | Grassland | 10° | 5° | 40 | 25 - 50 | -100 - -70 |
| M. canadense | Deciduous Broadleaf | 10° | 5° | 40 | 35 - 50 | -95 - -60 |

**Table S2:** Phenocam sites used in the analysis.

| **Site** | **Primary Vegetation** | **Matched Species** | **Latitude** | **Longitude** | **ROI Used** |
| --- | --- | --- | --- | --- | --- |
| marena | Grassland | R. hirta | 36.06434 | -97.21271 | 1000 |
| konza | Grassland | R. hirta | 39.08240 | -96.56030 | 1000 |
| uiefprairie | Grassland | R. hirta | 40.06462 | -88.19847 | 1000 |
| ninemileprairie | Grassland | R. hirta | 40.86802 | -96.82208 | 2000 |
| oakville | Grassland | R. hirta | 47.89933 | -97.31614 | 1000 |
| harvardfarmnorth | Grassland | R. hirta | 42.52047 | -72.18224 | 1000 |
| shiningrock | Deciduous Broadleaf | M. canadense | 35.39016 | -82.77497 | 3000 |
| harvard | Deciduous Broadleaf | M. canadense | 42.53780 | -72.17150 | 1000 |
| proctor | Deciduous Broadleaf | M. canadense | 44.52500 | -72.86600 | 1000 |
| willowcreek | Deciduous Broadleaf | M. canadense | 45.80599 | -90.07912 | 1000 |
| boundarywaters | Deciduous Broadleaf | M. canadense | 47.94670 | -91.49551 | 4000 |
| shalehillsczo | Deciduous Broadleaf | M. canadense | 40.66580 | -77.90410 | 2000 |

**Figure S1:**
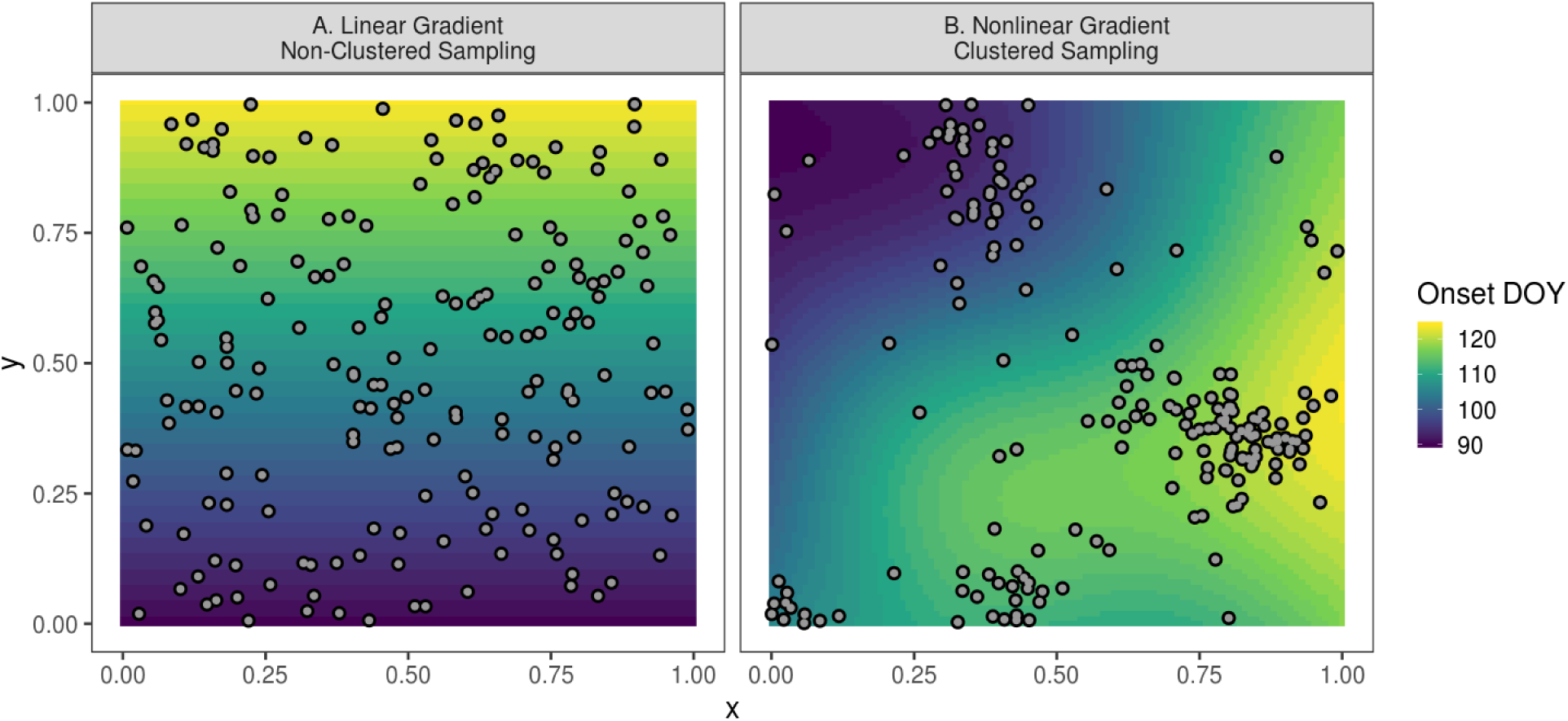
Example of simulated data. The color represents the onset, while points represent 200 sampled observations. Panel A depicts a linear spatial gradient which varies as a function of the y-axis and observations representing non-clustered sampling. Panel B depicts a nonlinear spatial gradient and observations representing clustered sampling.

**Figure S2:**
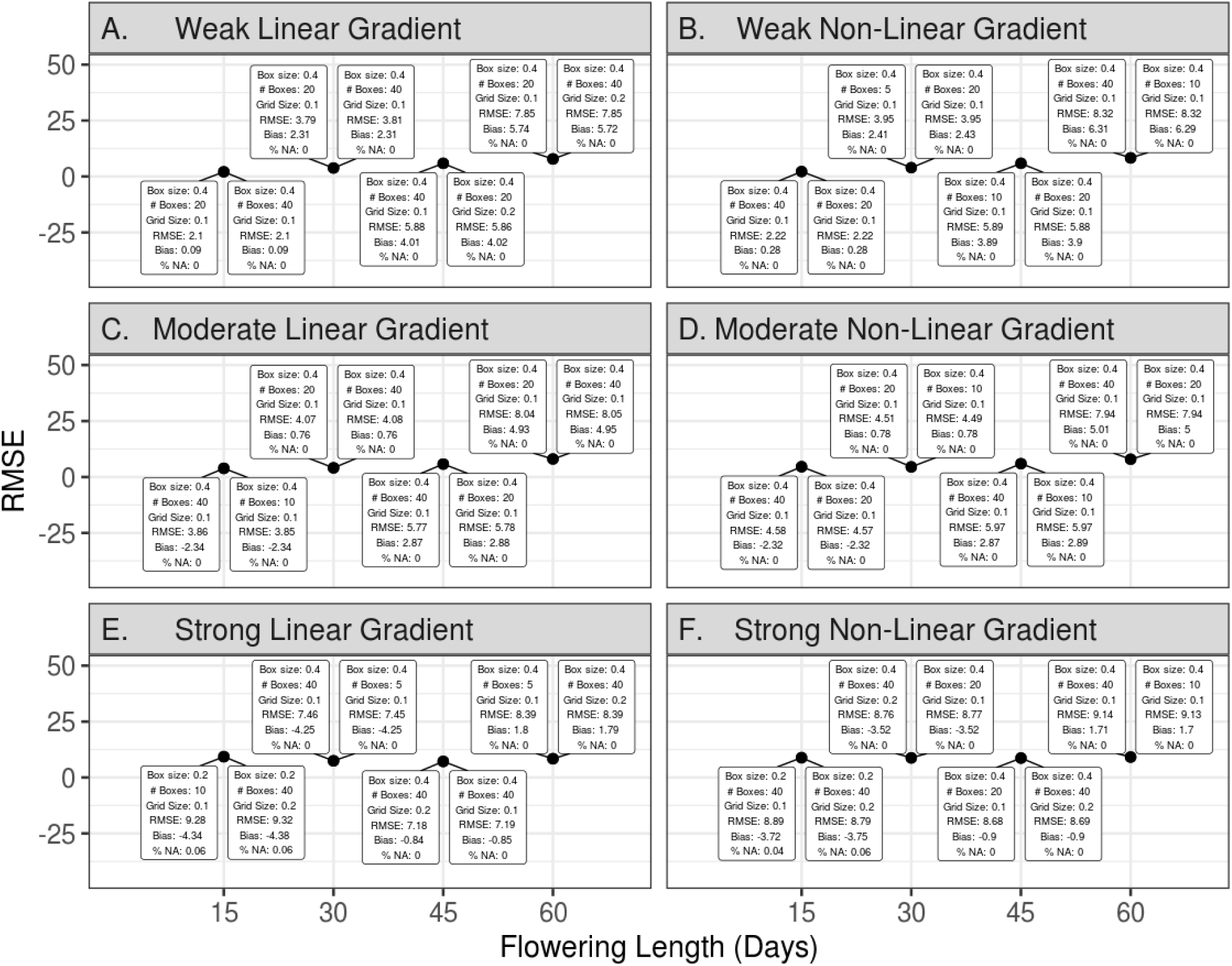
Weibull Grid parameters for the best performing models within each scenario. Points represent the RMSE of the Weibull Grid model for 6 types of flowering gradients (A-F) and 4 flowering season lengths (x-axis), and using a sample size of 300 based on clustered sampling. Text boxes indicate the two best performing parameter sets for that scenario as determined by lowest RMSE.

**Figure S3:**
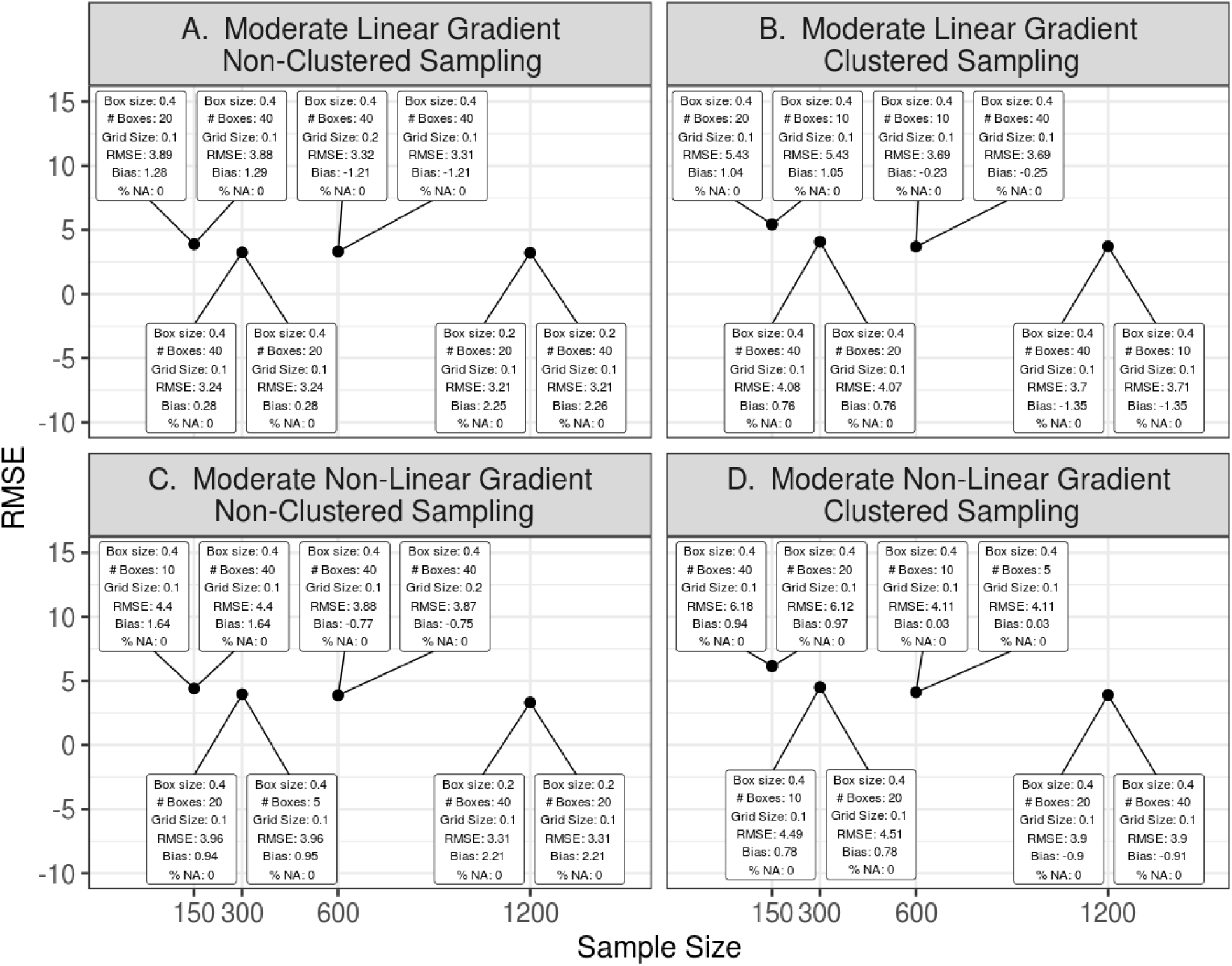
Weibull Grid parameters for the best performing model with in different sampling scenarios. Points represent the RMSE of the Weibull Grid model for either non-clustered (A-B) or clustered sampling (C-D). The underlying phenology is a moderate spatial gradient with a length of 30 days and spatial gradient linearity specified. Text boxes indicate the two best performing parameter sets for that scenario as determined by lowest RMSE.

**Figure S4:**
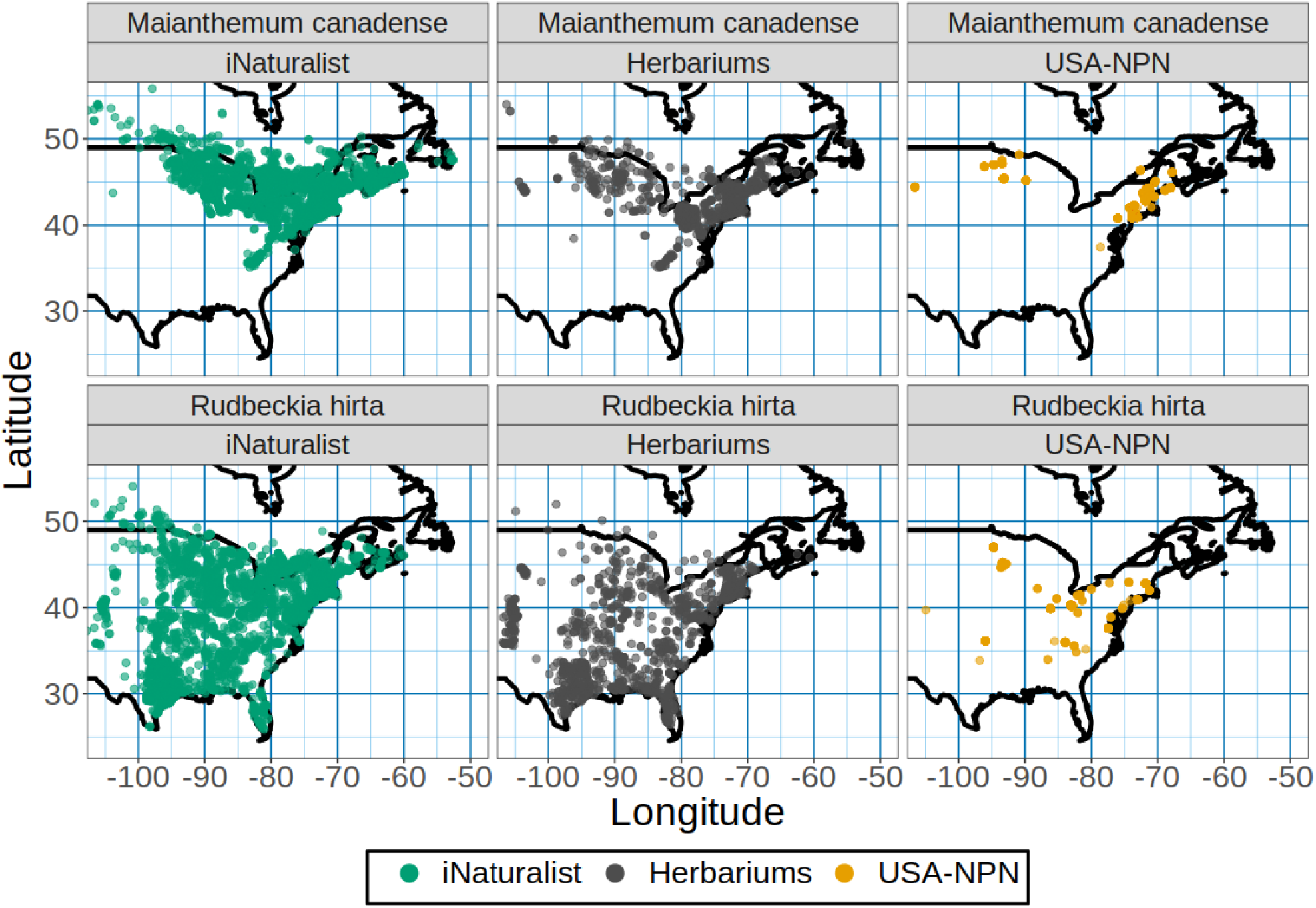
Locations of all data used in the analysis and discussion.

## LITERATURE CITED

Ault, T. R., M. D. Schwartz, R. Zurita-Milla, J. F. Weltzin, and J. L. Betancourt. 2015. Trends and Natural Variability of Spring Onset in the Coterminous United States as Evaluated by a New Gridded Dataset of Spring Indices. Journal of Climate 28: 8363–8378. https://doi.org/10.1175/JCLI-D-14-00736.1.

Baddeley, A., E. Rubak, and R. Turner. 2015. Spatial point patterns: methodology and applications with R. Chapman and Hall/CRC.

Barve, V. V, L. Brenskelle, D. Li, B. J. Stucky, N. V Barve, M. M. Hantak, B. S. Mclean, et al. Methods for broad-scale plant phenology assessments using citizen scientists’ photographs. bioRxiv: 754275. https://doi.org/10.1101/754275.

Brenskelle, L., B. J. Stucky, J. Deck, R. Walls, and R. P. Guralnick. 2019. Integrating herbarium specimen observations into global phenology data systems. Applications in Plant Sciences 7: e01231. https://doi.org/10.1002/aps3.1231.

Chuine, I., and J. Régnière. 2017. Process-Based Models of Phenology for Plants and Animals. Annual Review of Ecology, Evolution, and Systematics 48: 159–182. https://doi.org/10.1146/annurev-ecolsys-110316-022706.

Clark, J. S., J. Melillo, J. Mohan, and C. Salk. 2014. The seasonal timing of warming that controls onset of the growing season. Global Change Biology 20: 1136–1145. https://doi.org/10.1111/gcb.12420.

Flora of North America Editorial Committee. 1993. Flora of North America: Magnoliophyta: Liliidae: Liliales and Orchidales. Oxford University Press on Demand.

Daru, B. H., D. S. Park, R. B. Primack, C. G. Willis, D. S. Barrington, T. J. S. Whitfeld, T. G. Seidler, et al. 2018. Widespread sampling biases in herbaria revealed from large-scale digitization. New Phytologist 217: 939–955. https://doi.org/10.1111/nph.14855.

Dickinson, J. L., J. Shirk, D. Bonter, R. Bonney, R. L. Crain, J. Martin, T. Phillips, and K. Purcell. 2012. The current state of citizen science as a tool for ecological research and public engagement. Frontiers in Ecology and the Environment 10: 291–297. https://doi.org/10.1890/110236.

Dunnell, K. L., and S. E. Travers. 2011. Shifts in the flowering phenology of the northern Great Plains: Patterns over 100 years. American Journal of Botany 98: 935–945. https://doi.org/10.3732/ajb.1000363.

Elmendorf, S. C., T. M. Crimmins, K. L. Gerst, and J. F. Weltzin. 2019. Time to branch out? Application of hierarchical survival models in plant phenology. Agricultural and Forest Meteorology 279: 107694. https://doi.org/10.1016/j.agrformet.2019.107694.

Ettinger, A. K., S. Gee, and E. M. Wolkovich. 2018. Phenological sequences: how early-season events define those that follow. American Journal of Botany 105: 1771–1780. https://doi.org/10.1002/ajb2.1174.

Fink, D., W. M. Hochachka, B. Zuckerberg, D. W. Winkler, B. Shaby, M. A. Munson, G. Hooker, et al. 2010. Spatiotemporal exploratory models for broad-scale survey data. Ecological Applications 20: 2131–2147. https://doi.org/10.1890/09-1340.1.

Gazdic, M., and Q. Groom. 2019. iNaturalist is an Unexploited Source of Plant-Insect Interaction Data. Biodiversity Information Science and Standards, https://doi.org/10.3897/biss.3.37303.

GBIF.org. 2019. GBIF Occurrence Download. https://doi.org/10.15468/dl.zloi01.

Gerst, K. L., J. L. Kellermann, C. A. F. Enquist, A. H. Rosemartin, and E. G. Denny. 2016. Estimating the onset of spring from a complex phenology database: trade-offs across geographic scales. International Journal of Biometeorology 60: 391–400. https://doi.org/10.1007/s00484-015-1036-4.

Gräler, B., E. Pebesma, and G. Heuvelink. 2016. Spatio-Temporal Interpolation using gstat. The R Journal 8: 204–218.

Grolemund, G., and H. Wickham. 2011. Dates and Times Made Easy with lubridate. Journal of Statistical Software 40: 1–25.

Heberling, J. M., and B. L. Isaac. 2018. iNaturalist as a tool to expand the research value of museum specimens. Applications in Plant Sciences 6: e01193. https://doi.org/10.1002/aps3.1193.

Helenurm, K., and S. C. H. Barrett. 1987. The reproductive biology of boreal forest herbs. II. Phenology of flowering and fruiting. Canadian Journal of Botany 65: 2047–2056. https://doi.org/10.1139/b87-279.

Jelinski, D. E., and J. Wu. 1996. The modifiable areal unit problem and implications for landscape ecology. Landscape Ecology 11: 129–140. https://doi.org/10.1007/BF02447512.

de Keyzer, C. W., N. E. Rafferty, D. W. Inouye, and J. D. Thomson. 2017. Confounding effects of spatial variation on shifts in phenology. Global Change Biology 23: 1783–1791. https://doi.org/10.1111/gcb.13472.

Kosmala, M., A. Crall, R. Cheng, K. Hufkens, S. Henderson, and A. Richardson. 2016. Season Spotter: Using Citizen Science to Validate and Scale Plant Phenology from Near-Surface Remote Sensing. Remote Sensing 8: 726. https://doi.org/10.3390/rs8090726.

Kreyling, J., A. H. Schweiger, M. Bahn, P. Ineson, M. Migliavacca, T. Morel-Journel, J. R. Christiansen, et al. 2018. To replicate, or not to replicate – that is the question: how to tackle nonlinear responses in ecological experiments. Ecology Letters 21: 1629–1638. https://doi.org/10.1111/ele.13134.

Melaas, E. K., M. A. Friedl, and A. D. Richardson. 2016. Multiscale modeling of spring phenology across Deciduous Forests in the Eastern United States. Global Change Biology 22: 792–805. https://doi.org/10.1111/gcb.13122.

Melaas, E. K., D. Sulla-Menashe, and M. A. Friedl. 2018. Multidecadal Changes and Interannual Variation in Springtime Phenology of North American Temperate and Boreal Deciduous Forests. Geophysical Research Letters 45: 2679–2687. https://doi.org/10.1002/2017GL076933.

USA National Phenology Network. 2019. Plant and Animal Phenology Data. Data type: Status and Intensity. 01/01/2009-08/01/2019 for Region: 49.9375°, −66.4791667° (UR); 24.0625°, −125.0208333° (LL). USA-NPN, Tucson, Arizona, USA. Data set accessed 14 Sep 2019 at NPN, Tucson, Arizona, USA. Data set accessed 14 Sep 2019 at https://doi.org/10.5066/f78s4n1v.

Ogle, K., J. J. Barber, G. a. Barron-Gafford, L. P. Bentley, J. M. Young, T. E. Huxman, M. E. Loik, and D. T. Tissue. 2015. Quantifying ecological memory in plant and ecosystem processes. Ecology Letters 18: 221–235. https://doi.org/10.1111/ele.12399.

Parmesan, C., and G. Yohe. 2003. A globally coherent fingerprint of climate change impacts across natural systems. Nature 421: 37–42. https://doi.org/10.1038/nature01286.

Pearse, W. D., C. C. Davis, D. W. Inouye, R. B. Primack, and T. J. Davies. 2017. A statistical estimator for determining the limits of contemporary and historic phenology. Nature Ecology & Evolution 1: 1876–1882. https://doi.org/10.1038/s41559-017-0350-0.

Richardson, A. D., K. Hufkens, T. Milliman, D. M. Aubrecht, M. Chen, J. M. Gray, M. R. Johnston, et al. 2018. Tracking vegetation phenology across diverse North American biomes using PhenoCam imagery. Scientific Data 5: 1–24. https://doi.org/10.1038/sdata.2018.28.

Roberts, D. L., and A. R. Solow. 2003. When did the dodo become extinct? Nature 426: 245–245. https://doi.org/10.1038/426245a.

Scheffers, B. R., L. De Meester, T. C. L. Bridge, A. A. Hoffmann, J. M. Pandolfi, R. T. Corlett, S. H. M. Butchart, et al. 2016. The broad footprint of climate change from genes to biomes to people. Science 354: aaf7671. https://doi.org/10.1126/science.aaf7671.

Stucky, B. J., R. Guralnick, J. Deck, E. G. Denny, K. Bolmgren, and R. Walls. 2018. The Plant Phenology Ontology: A New Informatics Resource for Large-Scale Integration of Plant Phenology Data. Frontiers in Plant Science 9. https://doi.org/10.3389/fpls.2018.00517.

Taylor, S. D. 2019. Estimating flowering transition dates from status-based phenological observations: a test of methods. PeerJ 7: e7720. https://doi.org/10.7717/peerj.7720.

R Core Team. 2017. R: a language and environment for statistical computing.

Theobald, E. J., I. Breckheimer, and J. HilleRisLambers. 2017. Climate drives phenological reassembly of a mountain wildflower meadow community. Ecology 98: 2799–2812. https://doi.org/10.1002/ecy.1996.

Weston, S., and R. Calaway. 2019. Getting Started with doParallel and foreach.

Wickham, H. 2016. ggplot2: Elegant Graphics for Data Analysis. Springer-Verlag New York.

Wickham, H. 2011. testthat: Get Started with Testing. The R Journal 3: 5–10.

Wickham, H., R. Francois, L. Henry, and K. Müller. 2017. dplyr: A Grammar of Data Manipulation.

Wickham, H., and L. Henry. 2018. tidyr: Easily Tidy Data with ‘spread()’ and ‘gather()’ Functions.

Willis, C. G., E. R. Ellwood, R. B. Primack, C. C. Davis, K. D. Pearson, A. S. Gallinat, J. M. Yost, et al. 2017. Old Plants, New Tricks: Phenological Research Using Herbarium Specimens. Trends in Ecology & Evolution 32: 531–546. https://doi.org/10.1016/j.tree.2017.03.015.

Wolkovich, E. M., and A. K. Ettinger. 2014. Back to the future for plant phenology research. New Phytologist 203: 1021–1024. https://doi.org/10.1111/nph.12957.

